# miR-6850 Drives Phenotypic Changes and Signaling in High Grade Serous Ovarian Cancer

**DOI:** 10.1101/2025.07.30.667620

**Authors:** Kamil Filipek, Daniela Pollutri, Ivana Kurelac, Giuseppe Gasparre, Marianna Penzo

## Abstract

MicroRNAs (miRNAs) are key post-transcriptional regulators of gene expression, and their dysregulation is closely linked to cancer development. Ovarian cancer (OC), particularly the high-grade serous ovarian carcinoma (HGSOC) subtype, is the most lethal gynecological malignancy, primarily due to late-stage diagnosis and limited treatment options. Among the miRNAs encoded at the often amplified 8q24.3 region, miR-6850 has emerged as a potential candidate target owing to its genomic positioning inside this hotspot and its unexpectedly low expression in HGSOC tissues and cell lines. In silico investigations indicated that, despite the gain in MIR6850 copy number, its mature products, miR-6850-5p and miR-6850-3p, were expressed at low levels; notably, *MIR6850* gene amplification was associated with enhanced disease-specific survival. Functional studies revealed that ectopic production of both isoforms in SKOV-3 and NIH:OVCAR3 cells inhibited proliferation, compromised clonogenic capacity, and disturbed cell cycle progression. Moreover, miR-6850 altered cell phenotype by facilitating mesenchymal-to-epithelial transition (MET), as shown by the overexpression of E-cadherin and β-catenin and the downregulation of Slug and Vimentin. It also regulated cell adhesion and migration while reducing global protein synthesis via the downregulation of the PI3K/Akt/mTOR pathway. Our results together identify miR-6850 as a tumor-suppressive miRNA in HGSOC, demonstrating its diverse anti-oncogenic actions and underscoring its potential as a prognostic biomarker and therapeutic target in ovarian cancer.

## 1. Introduction

OC is currently one of the most worrying medical-biological problems as it remains the most fatal form of malignancy in the gynecological field [1], being the eighth most common cancer in women worldwide [2]. In 2022, Globocan reported 324,398 new OC diagnoses and 206,839 OC-related deaths [3]. The survival rate of patients 5 years after diagnosis is dramatically low, around 40%. This is attributable to the fact that the tumor is diagnosed when it is already in an advanced stage (FIGO III/IV), due to the late onset of symptoms [1] and to the limited availability of targeted therapies [4]. These premises underline the need to delve deeper into the molecular pathogenesis of OC, to identify novel players or pathways that could be used as biomarkers, for risk stratification, or as druggable targets for OC treatment.

miRNAs are a class of small (19-22 nucleotides) non-coding RNA molecules distributed throughout the genome, which contribute to regulating gene expression [5], thus playing an important role in many physiological cellular processes [6]. Their de-regulated expression is known to be involved in different pathological conditions, especially in tumors, including OC [7–9], and can be correlated with a specific clinical outcome for patients [10].

Several studies have demonstrated that miRNAs are linked to the development of OC. For instance, miR-216a promoted OC progression by the downregulation of the tumor suppressor PTEN [7]. Downregulation of miR-451a fostered OC proliferation by targeting the Ras-ERK pathway [8], whereas miR-149-3p stimulated the cisplatin resistance and promoted tumorigenesis through the downregulation of CDKN1A and TIMP2 proteins [9]. miR-6850 is a novel disease-related miRNA. The upregulated expression of miR-6850-5p has been shown in colorectal cancer [11], endometrial cancer cells [12], in the serum of patients with type 2 diabetes mellitus [13], following exposure to gamma radiation [14], and in cerebrospinal fluids of patients with febrile seizures [15]. In contrast, downregulation of miR-6850-5p expression has been described in the serum of patients with centrally mediated abdominal pain syndrome (CAPS), for whom it has been proposed as a promising diagnostic biomarker [16]. However, the expression and biological role of miR-6850 in HGSOC have never been investigated.

In this study, we found that miR-6850-5p and miR-6850-3p are expressed at low levels in ovarian cancer tissues and cell lines, despite frequent genomic amplification of the *MIR6850*. Notably, this amplification is associated with improved disease-specific survival. Functional assays further revealed that enforced expression of either strand leads to a less aggressive cellular phenotype, supporting a tumor-suppressive role for miR-6850 in HGSOC.

## 2. Materials and Methods

### 2.1. Cell cultures and transfection

Ovarian cell lines SKOV-3 and NIH:OVCAR3 were purchased from H□lzel Diagnostika (Cologne, Germany), OVSAHO from JCRB Cell Bank and Sekisui XenoTech, LLC, (Kansas City, KS, USA), CAOV-3 (#ATCC-HTB-75) and OV-90 (#ATCC-CRL-3585) from ATCC. OC314 and, human ovarian surface epithelial (HOSE) were obtained from a third part. Cell line authentication (Suppl. Figure 1) was performed as previously described [17, 18] either by TP53 Sanger sequencing to identify the characteristic mutations or by AMpFISTR Identifiler (Applied Biosystems). The OC cell lines were cultured in RPMI-1640 medium (SIAL-RPMI-XA, Sial, Rome, Italy) supplemented with 10% FBS (YOURSIAL-FBS-SA, Sial), 100 U/ml penicillin, and 100 µg/ml streptomycin (SIAL-PEN/STREP, Sial), and 2 mM L-glutamine (SIAL-Lglu, Sial). HOSE cell line was cultured in ovarian epithelial cell medium supplemented with ovarian cell growth and penicillin-streptomycin solution (Innoport, Spain). Cultures were maintained in a humidified atmosphere with 5% CO□ at 37 °C.

SKOV-3 and NIH:OVCAR3 cell lines were transfected with miRNA mimics for miR-6850-5p (Assay ID: MC28151, ThermoFisher Scientific, Waltham, MA, USA), miR-6850-3p (Assay ID: MC29832, ThermoFisher Scientific) using Lipofectamine RNAiMAX (Invitrogen), following the manufacturer’s protocol. Cells at approximately 70% confluence were transfected with 40 pmol of each mimic. As a control, cells were transfected with Opti-MEM and RNAiMAX and are hereafter referred to as “vehicle”.

### 2.2 Total RNA extraction, reverse transcription and real time PCR

RNA extraction was performed utilizing the PureZOL reagent (Bio-Rad, Hercules, CA, USA), according to the manufacturer’s instructions. A 10 ng and 500 ng of total RNA was retrotranscribed using TaqMan MicroRNA Reverse Transcription Kit (ThermoFisher Scientific) and iScript cDNA Synthesis Kit (Bio-Rad), respectively. The specific primers for U6 snRNA (Assay ID: 001973 ThermoFisher Scientific), miR-6850-5p (Assay ID: 466633_mat, ThermoFisher Scientific), and miR-6850-3p (Assay ID:466668_mat, ThermoFisher Scientific) were added to the RNA templates, followed by incubation at 85 °C for 5 minutes, 60°C for 5 minutes, and cooling on ice. The RT for miRNA was performed using these conditions: 16 °C for 30 minutes, 42°C for 30 minutes, and 85°C for 5 minutes, followed by cooling at 4°C. The RT for mRNA was performed using these conditions: 25°C for 5 minutes, 46°C for 25 minutes, and 95°C for 1 minute. The cDNA was diluted with RNAse-free water in 1:5 ratio.

Real-time PCR for miRNA was performed using TaqMan Universal Master Mix (ThermoFisher Scientific), probes for U6 snRNA (internal control), miR-6850-5p, and miR-6850-3p (ThermoFisher Scientific). Thermal conditions: 95°C for 10 minutes, 40 cycles at 95 °C for 15 seconds, and 60°C for 1 minute.

Real-time PCR for miR-6850 potential targets was performed using SsoAdvanced Universal SYBR Green Supermix (Bio-Rad) and specific primers provided in table 1 below. TBP (TATA binding protein) was used as an iternal control.Thermal conditions: 95°C for 10 minutes, 40 cycles at 95°C for 15 seconds, and 60°C for 1 minute. The fold change was calculated according to the relative quantification method (2^-ΔΔCt^).

**Table 1.**
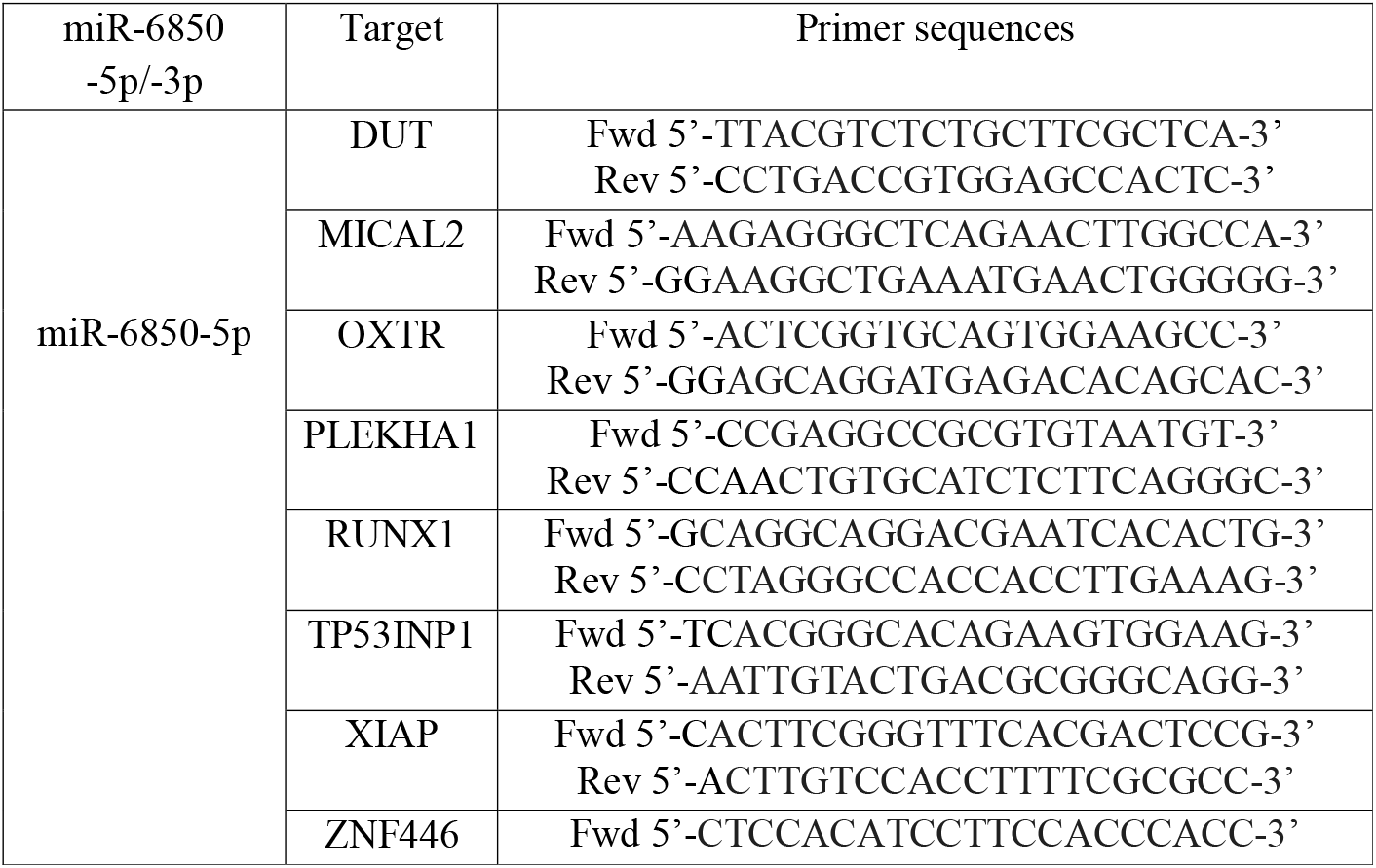
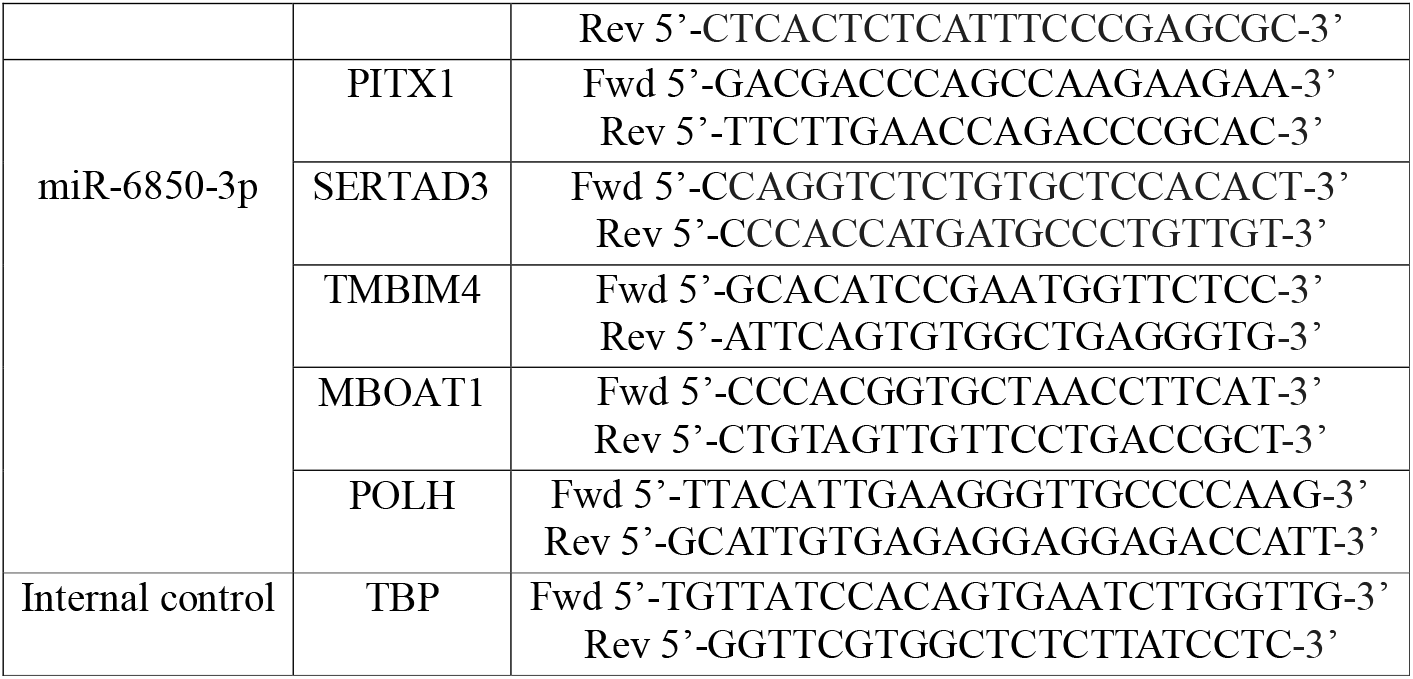
Primer sequences for miR-6850 potential targets.

### 2.5 Protein extraction and immunoblotting

Whole cell protein extracts were prepared in RIPA buffer and quantified using Bio-Rad Protein Assay Dye Reagent Concentrate. 20 µg of proteins were resolved by SDS-PAGE electrophoresis in 12% polyacrylamide gels and transferred to nitrocellulose membranes (Amersham, Merck, Darmstadt, Germany) using a Mini Trans-Blot Cell Module (Bio-Rad). After transfer, the membranes were blocked in 5% non-fat dry milk in TBST buffer (50 mM Tris-HCl pH 7.4, 150 mM NaCl, 0.1% Tween-20) for 1 h at RT with gentle shaking, and incubated overnight at 4 °C with indicated primary antibodies diluted in TBST with 3% BSA. After washing, a 1 h incubation with the relevant secondary antibody was performed, and detection was carried out with chemiluminescent signals were developed using Westar Antares (Cyanagen, Bologna, Italy) using ChemiDoc™ XP+ (Bio-Rad), and Image Lab 6.0 software (Bio-Rad). The list of used antibodies was described in Table S1.

### 2.6 Cell proliferation assay

SKOV-3 and NIH:OVCAR3 7500 cells per well were seeded into a 6-well plate. After 2 days, the cells were transfected with miR-6850-5p, miR-6850-3p, or vehicle. SKOV-3 and NIH:OVCAR3 growth levels were assessed every 24 hours over a total duration of 7 days. For cell counting, cells were mixed with 0.1% erythrosine in a 1:1 volume ratio, and the counting mix was loaded into a counting chamber and analyzed using Countess II FL (ThermoFisher Scientific). All counts were performed in triplicate.

### 2.7 Cell cycle analysis

To assess the cell cycle 72 hours post-transfection, cells were resuspended in DPBS at a concentration of 10□ cells/ml. After fixing with 2.5 volumes of 100% pure ethanol at RT with moderate agitation, the cells were incubated overnight at −20 °C. The next day, the cells were pelleted, rinsed with DPBS and treated with propidium iodide (10 µg/ml) (Merck) and RNase A (100 µg/ml) (ThermoFisher Scientific) at 37 °C for 20 minutes. Following incubation, stained cells were resuspended in 500 µl of DPBS and stored in the dark on ice. A fluorescence-negative control (PI-free cells) was also prepared. The CytoFLEX Flow Cytometer and Kaluza Analysis Software (both from Beckman Coulter, Brea, CA, USA) were used to acquire the cell cycle and to analyze the cell cycle, respectively.

### 2.8 Clonogenic assay

SKOV-3 and NIH:OVCAR3 cells were transfected with miR-6850-5p, miR-6850-3p or vehicle as previously described. 24 hours post-transfection, 100 cells/well of SKOV-3 and 400 cells/well of NIH:OVCAR3 were seeded into a 6-well plate in triplicate. The colony number was evaluated after 12 days of culture. Briefly, cells were washed twice with DPBS and fixed overnight in 4% formalin at 4 °C. The next day, the cells were stained with a 0.5% crystal violet solution in 25% methanol for 30 min. Cells were then washed 3 times in DPBS and colonies were counted.

### 2.9 Adhesion assay

A 96-well plate was used to assess SKOV-3 and NIH:OVCAR3 cell adhesion 3 days post-transfection with miR-6850-5p, miR-6850-3p or vehicle. Wells were coated with Laminin-1 (10 μg/ml), Fibronectin (20 μg/ml), Gelatin (0.2%), Collagen I (10 μg/cm2), and Poly-L-Lysine (100 μg/ml) at 37 °C for 1 hour. Control wells were left uncoated. Post-coating, wells were washed twice with 0.1% BSA in RPMI-1640 and blocked with 0.5% BSA at 37 °C for 1 hour. Wells were buffer washed three times. Transfected SKOV-3 and NIH:OVCAR3 cells were resuspended at the concentration of 4 ×10^5^ cells/ml, and 50 µl of cell suspension per well were seeded in each well, then incubated at 37 °C, 5% CO□ for 30 minutes. After, the plate was shaken at 2,000 RPM for 15 seconds and the wells were washed three times with a washing buffer to eliminate non-adherent cells. Adherent cells were fixed with 4% paraformaldehyde for 15’ at RT, washed twice with washing buffer, and stained with Crystal Violet (5 mg/ml in 2% ethanol) for 10 minutes. The wells were washed with water and dried completely by inversting the plates. After solubilizing the dye with 2% SDS, the plates were incubated at room temperature for 30 minutes. Absorbance at 550 nm was measured in a Spark plate reader (Tecan, Männedorf, Switzerland).

### 2.10 Scratch assay

SKOV-3 and NIH:OVCAR3 cells were transfected with miR-6850-5p, miR-6850-3p or vehicle as described above. After 2 days, 50,000 cells/well were seeded into 96-well IncuCyte^®^ ImageLock microplate (Essen BioScience, Ann Arbor, MI, USA) wells pre-coated with 0.2% gelatin and cultured for one day. Then, confluent cells were mechanically scratched using a 96-pin IncuCyte WoundMaker Tool (Essen BioScience), creating 700-800 µm wide wounds. The detached cells were removed by washing the wells twice with DPBS, and the wells were filled with 100 µl of culture medium. The plate was inserted into the IncuCyte S3 system (Sartorius, Göttingen, Germany) and images were captured every 2 h for SKOV-3 and 3 h for NIH:OVCAR3 up to 48 h after scratching. The relative wound confluence (%) was quantified by the Cell Migration Software Application Module (Essen BioScience).

### 2.11 Accession of the public database

The TCGA PanCancer Atlas database deposited in cBioPortal was used to acquire the *MIR-6850* amplification percentage, gene copy number and RPL8 mRNA expression values in ovarian serous cystadenocarcinoma tissues. The TCGA database and NCBI SRA (Sequence Read Archive) deposited in DIANA-microRNA Tissue Expression Database (miTED) were used to acquire the *MIR-6850, MIR-6849, MIR-661, MIR-937, MIR-939* expression values in ovarian serous cystadenocarcinoma tissues and cell lines, respectively.

### 2.12 Survival analysis

The HGSOC patients were divided into a genomic *MIR-6850* altered group and a *MIR-6850* unaltered group. The disease-specific survival was calculated by cBioPortal [19]. The hazard ratio with 95% confidence intervals and p-value were calculated and displayed.

### 2.13 In Silico prediction of miR-6850 target genes

Putative target genes of miR-6850-5p and miR-6850-3p were predicted using miRPathDB [20], DIANA-microT [21], miRTarBase [22], miRWalk [23], miRDB [24], and TargetScan (v8.0) [25] databases. Only targets identified by at least four databases were considered reliable and included in subsequent analyses.

### 2.14 Statistical analysis

All experiments were repeated at least three times. Values are presented as a mean ± SD. Statistical differences between two groups were assessed by two-tailed Student’s t-test. For correlation analysis, Pearson’s and Spearman correlation were used for calculating *r* and *p* values. Significant differences are indicated as follows: \**p*<0.05; \*\**p*<0.01; \*\*\**p*<0.001; \*\*\*\**p*<0.0001. GraphPad Prism 10.5.0 (GRAPH PAD Software Inc., La Jolla, CA, USA) was used for statistical analysis and plotting the graphs.

## 3. Results

### 3.1 MIR6850 genomic localization and status

Our lab has a longstanding interest in ribosome biology and the role of ribosomal genes in cancer. While investigating *RPL8*, a ribosomal protein gene frequently amplified in HGSOC, we identified miR-6850, a previously uncharacterized miRNA embedded within its first intron. This serendipitous observation prompted us to explore whether miR-6850 might contribute to OC pathogenesis, particulary given its genomic location in the 8q24.3 region – an area known for instability and recurrent amplification in HGSOC [26, 27]. To investigate the potential involvement of miR-6850 in HGSOC, we first examined its genomic organization. miR-6850 is transcribed from a single locus (*MIR6850*, Gene ID: 102465978), located on the reverse strand within intron 1 of the *RPL8* gene (Fig. 1A). The *MIR6850* gene gives rise to two mature isoforms: the 22bp long miR-6850-5p and the 20bp long miR-6850-3p, both processed from the same precursor.

**Fig. 1.**
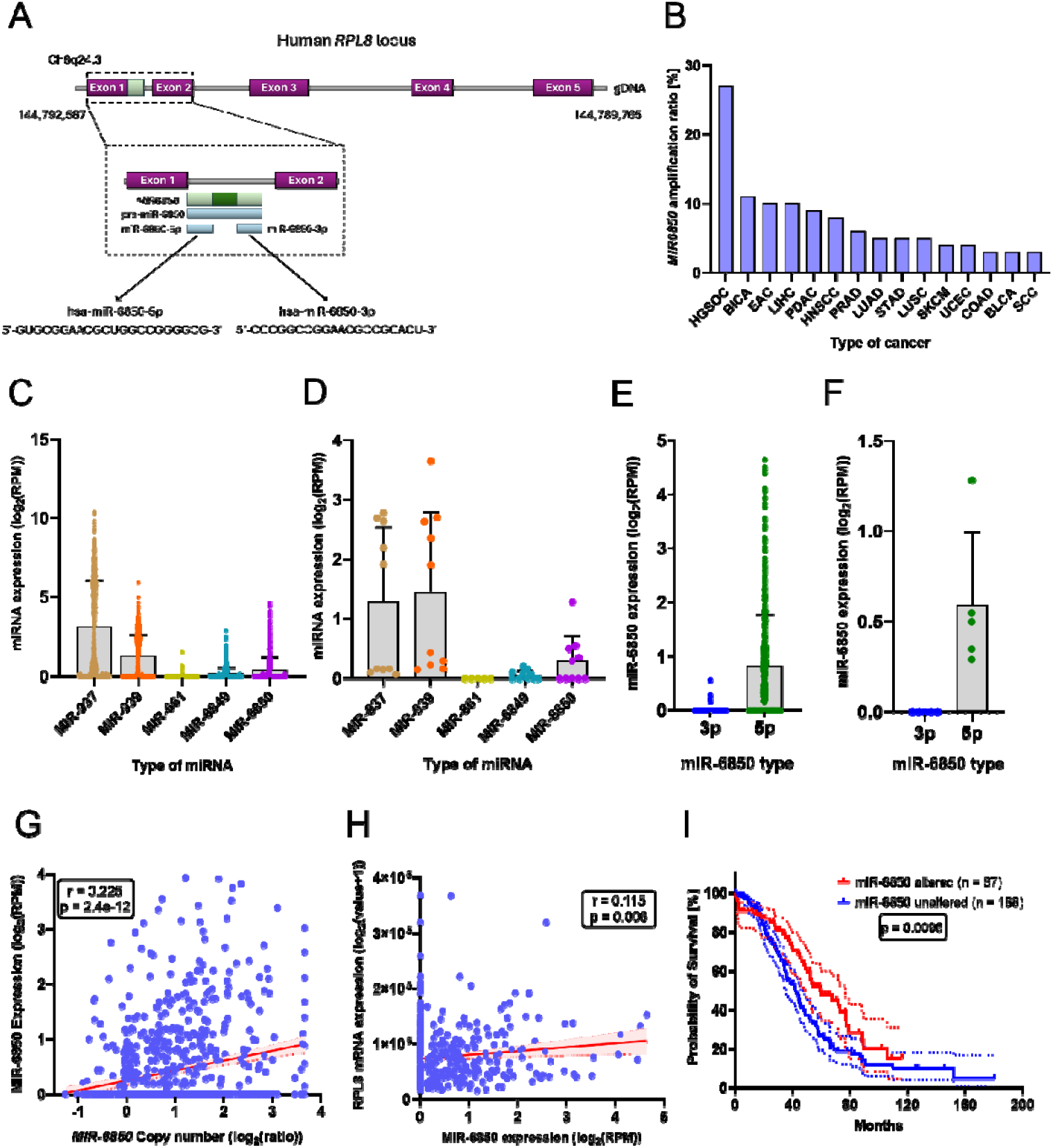
miR-6850 amplification and expression in ovarian cancer lines and clinical samples. (A) Schematic diagram of genomic localization of *MIR6850* gene within *RPL8* locus. (B) Genomic alteration of *MIR6850* across human cancers retrieved from TCGA PanCancer Atlas dataset deposited in cBioPortal. The expression values of miR-937, miR-939, miR-661, miR-6849, miR-6850 in ovarian cystadenocarcinoma samples from TCGA database (C) and cell lines from NCBI SRA database (D). The expression values of miR-6850-3p and miR-6850-5p in ovarian cystadenocarcinoma samples from TCGA database (E) and cell lines from NCBI SRA database (F). The bars indicate the median expression of each miRNA. The correlations between (G) *MIR6850* copy number and its expression and (H) miR-6850 expression and RPL8 mRNA expression using the TCGA database. The *r* and *p* values are displayed on the graphs. (I) Kaplan-Meier curve of disease-specific survival of patients based on the miR-6850 genetic status in TCGA cohort (n = 275). The number of patients as well as the *p-*value are presented in the graph.

To understand the clinical relevance of this locus, we next assessed the amplification frequency and expression profile of *MIR6850* in HGSOC patient samples and cell lines. Using the TCGA PanCancer Atlas dataset (via cBioPortal) [19], we observed that *MIR6850* is amplified in 27% of HGSOC samples – representing the highest amplification frequency among all human cancers considered (Fig. 1B). To gain insight into the broader expression context of this locus, we examined five microRNAs located on 8q24.3—miR-937, miR-939, miR-661, miR-6849, and miR-6850—in HGSOC tumor samples from the TCGA database. Among these, miR-937 and miR-939 showed the highest expression levels, while miR-6850 was one of the least expressed miRNAs in this region (Fig.1C). Consistent with this observation, analysis of RNA-sequencing data from HGSOC cell lines, retrieved from the NCBI Sequence Read Archive via the DIANA-microRNA Tissue Expression Database (miTED) [28], confirmed the relatively low abundance of miR-6850, which was intermediate compared to the higher expression of miR-937 and miR-939, and lower expression of miR-661 and miR-6849 (Fig.1D). Notably, across tumor samples (Fig.1E) and cell lines (Fig.1F) alike, miR-6850-5p was expressed at higher levels than miR-6850-3p.

To assess genomic influences on MIR6850 expression in HGSOC, we examined the correlation between its DNA copy number and RNA levels. This analysis revealed a significant positive correlation (r = 0.225, p = 2.4e-12), indicating that copy number gains may contribute to higher MIR6850 expression (Fig. 1G). We also evaluated the relationship between MIR6850 and its host gene RPL8, identifying a weak yet statistically significant positive correlation in their expression levels (r = 0.115, p = 0.006) (Fig. 1H).

Finally, we evaluated the clinical relevance of miR-6850 expression using Kaplan– Meier survival analysis in the TCGA HGSOC cohort (n = 275). Strikingly, higher miR-6850 amplification levels were significantly associated with improved disease-specific survival (p = 0.0096) (Fig.1I), suggesting a potential role for miR-6850 as a favorable prognostic biomarker in HGSOC.

### 3.2 miR-6850 regulates cell proliferation

These *in silico* results prompted us to advance our understanding of the role and underlying mechanism of miR-6850 in HGSOC. We first evaluated the basal expression of its mature strands, miR-6850-5p and miR-6850-3p, across a panel of serous ovarian carcinoma (SOC) cell lines (SKOV-3, OVSAHO, OV90, CAOV3, NIH:OVCAR3, OC314) and compared it to the human ovarian surface epithelial (HOSE) cell line, representing non-transformed ovarian epithelium. This initial analysis was essential for identifying suitable models to interrogate the functional impact of miR-6850 in distict cellular context. Consistent with our previous findings, miR-6850 expression was markedly reduced in all examined HGSOC cell lines relative to HOSE (Fig. 2A). The miR-6850-5p was consistently more abundant than miR-6850-3p across cancer cell lines; this strand bias was not evident in HOSE, suggesting subtype-specific regulation.

**Fig. 2.**
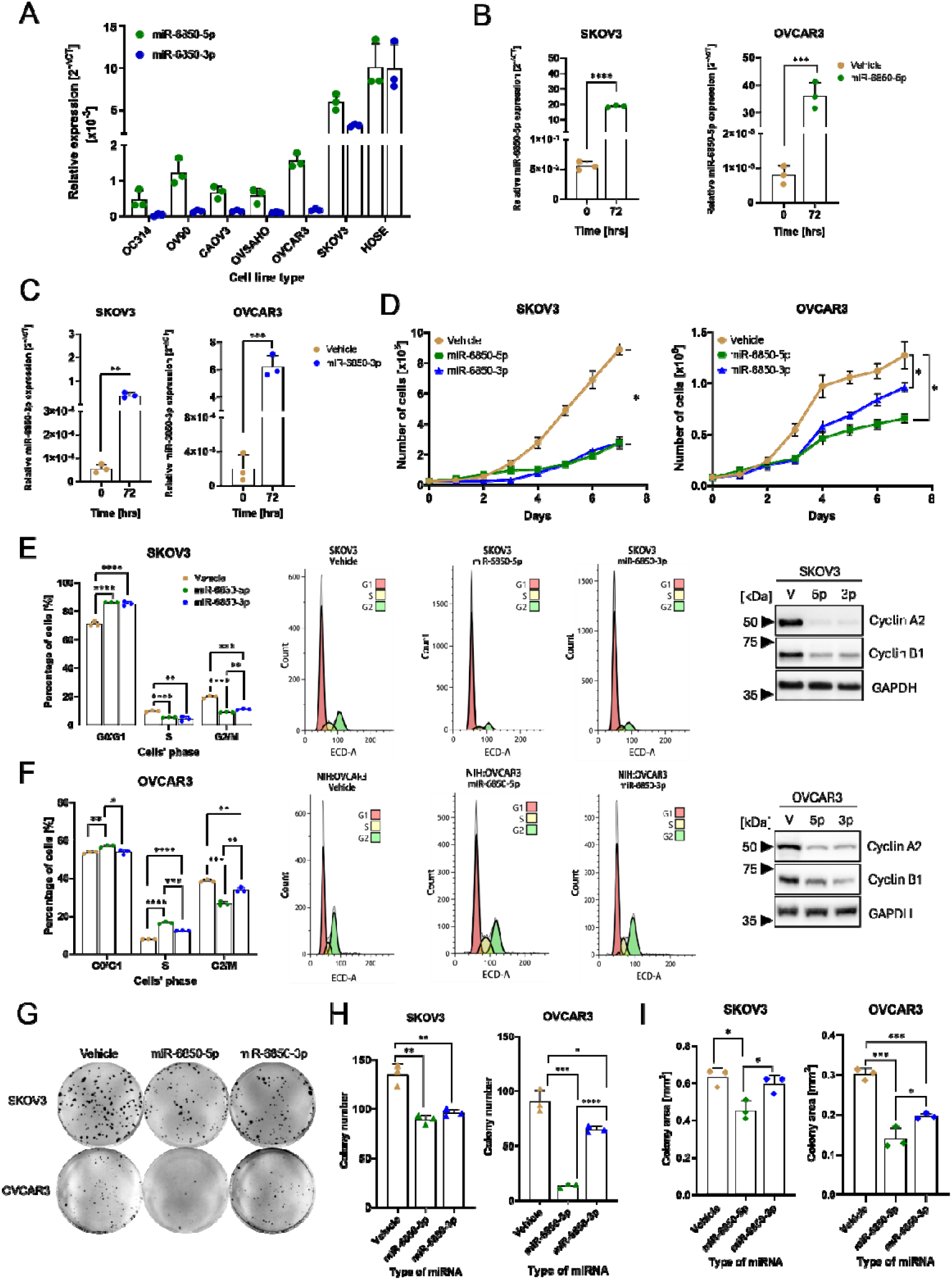
Functional impact of miR-6850 on cell proliferation.. (A) Basal expression of miR-6850-5p and miR-6850-3p in a panel of ovarian cell lines. Expression of miR-6850-5p (B) and miR-6850-3p (C) in transfected miRNA mimics SKOV-3 and NIH:OVCAR3 cell lines. (D) Growth rate up to 7 days of SKOV-3 and NIH:OVCAR3 cells after transfection with miR-6850-5p (green line), miR-6850-3p (blue line), and control cells (brown line). Cell cycle distribution and cyclins expression in (E) SKOV-3 and (F) NIH:OVCAR3 overexpressing miR-6850 and control cells (vehicle). The left panel shows the quantification graph of cells in different phases of cell cycle in control and miR-6850-5p/-3p transfected cells. The central panel presents the representative images of SKOV-3 and NIH:OVCAR3 cell cycle analysis acquired from Kaluza Software. The right panel shows the expression of Cyclin A2 and Cyclin B1 in both investigated cell lines at indicated time points. GAPDH was used as a loading control. (G-I) Clonogenic potential of SKOV-3 and NIH:OVCAR3 cells overexpressing miR-6850-5p, miR-6850-3p, and control cells. (G) Representative images of one biological replicate of clonogenic assay for SKOV-3 and NIH:OVCAR3. Quantification of (H) colony number and (I) average colony area reported in mm^2^. All experiments were conducted in three biological replicates and all data are shown as mean±SD. *p<0.05; **p<0.01; ***p<0.001, ****p<0.0001 determined by unpaired Student’s t-test.

For subsequent studies, we selected the SKOV-3 and NIH:OVCAR3 cell lines based on their known distinct molecular and phenotypic characteristics. SKOV-3 cells exhibit a partial EMT phenotype, characterized by low E-cadherin and high vimentin expression [29] and displayed relatively higher basal levels of miR-6850 (Fig.2A). Additionally, SKOV-3 harbors gain-of-function mutation in *PIK3CA*, leading to constitutive activation of the PI3K/Akt/mTOR pathway [30]. In contrast, NIH:OVCAR3 cells display an epithelial phenotype with robust E-cadherin expression [29], retain wild-type *PIK3CA* and *PTEN* [30], and expressed lower levels of miR-6850 (Fig.2A). The use of these two models enabled us to investigate miR-6850’s functional effects in both mesenchymal-like and epithelial cellular contexts.

To investigate the role of *MIR6850* in HGSOC progression, we overexpressed both miR-6850-5p and miR-6850-3p in SKOV-3 and NIH:OVCAR3 cell lines using miRNA mimics. The relative expression was quantified using qRT-PCR. The qRT-PCR analysis revealed that cells transfected with miR-6850-5p (Fig.2B) and miR-6850-3p (Fig.2C) mimics demonstrated an increase in expression over the observed period, compared to the control cells. We hence assessed the effect of miR-6850 on different cancer-relevant phenotypic outcomes in SKOV-3 and NIH:OVCAR3 cells, over a period of seven consecutive days. The results indicated a significant growth reduction in both SKOV-3 and NIH:OVCAR3 cells overexpressing both miR-6850-5p and miR-6850-3p (p<0.05) compared to the control group, even if with a variable degree in the different cellular contexts and, in NIH:OVCAR3 cells, with the different mimics (Fig.2D).

We therefore verified the cell cycle. In both analyzed cell lines, we observed significant changes in phase distribution three days post-transfection. In SKOV-3, overexpression of miR-6850-5p and miR-6850-3p led to a significant increase in G0/G1 phase cells and a decrease in both S and G2/M phase cells compared to control cells (Fig.2E left). On the other hand, in NIH:OVCAR3, we noticed a significant increase of cells in S phase following a decrease in G2/M phase (Fig.2F left). In the attempt to explain the observed differences in cell cycle phase distribution, we examined cyclin A2 and cyclin B1 expression. Indeed, in both SKOV-3 and NIH:OVCAR3 cells we found a marked decrease in protein level of both cyclin A2 and cyclin B1 at the third day after transfection with miR-6850-5p and miR-6850-3p (Fig.2E and 2F right). The effect of the −3p and −5p miRNAs was more repressive towards cyclin A2 rather than cyclin B1.

Finally, we assessed the impact of miR-6850-5p and miR-6850-3p overexpression on the clonogenic potential of OC cells by comparing the number and size of colonies formed, compared to control cells. Overall, SKOV-3 cells showed a higher capacity for single colony formation compared to NIH:OVCAR3 cells, as illustrated in Figure 2G. In detail, transfection of SKOV-3 and NIH:OVCAR3 cells with miR-6850-5p/-3p mimics resulted in a decrease in colony-forming ability, and the strongest inhibitory effect was described for NIH:OVCAR3 overexpressing miR-6850-5p (Fig.2H). In terms of colony area, the colonies generated by the miR-6850-5p transfected SKOV-3 and NIH:OVCAR3 cells exhibited a reduced size, averaging 28% and 53% smaller in area compared to the control group, respectively. For miR-6850-3p generated colonies, their size was reduced by 6% and 35% for SKOV3 and NIH:OVCAR3 cells, respectively (Fig.2I).

Taken together, these results suggest a tumor-suppressive role of miR-6850 in HGSOC, potentially acting through regulation of cell cycle and proliferation-associated pathways.

### 3.3 miR-6850 regulates cell adhesion, migration, and EMT process

We next investigated the effect of miR-6850-5p and miR-6850-3p on the adhesion potential of SKOV-3 and NIH:OVCAR3 cells, exploring different adhesive substrates: laminin-1 (Lmn-1), fibronectin (Fbn), gelatin (Gel), collagen I (Col 1), and poly-L-lysine (P-Lys). Figure 3A illustrates that cells overexpressing miR-6850-5p exhibited increased adherence, while those overexpressing miR-6850-3p demonstrated reduced adherence to all substrates and noncoated wells in comparison to control cells. In SKOV-3 cells overexpressing −5p, we observed the strongest effects on adhesion to gelatin and poly-L-lysine, with adhesive potential increased by approximately 185% and 263%, respectively, compared to the control group. On the other hand, for SKOV-3 overexpressing miR-6850-3p the adhesion to fibronectin and gelatin decreased by approximately 57% and 50%, respectively, compared to control (Fig.3A left). In contrast, NIH:OVCAR3 had generally higher adhesive potential compared to SKOV-3 cells. Nevertheless, we noted some changes in the adhesive potential of NIH:OVCAR3 cell upon −3p and −5p overexpression, albeit significant only for gelatin (Fig.3A lower panel).

**Fig. 3.**
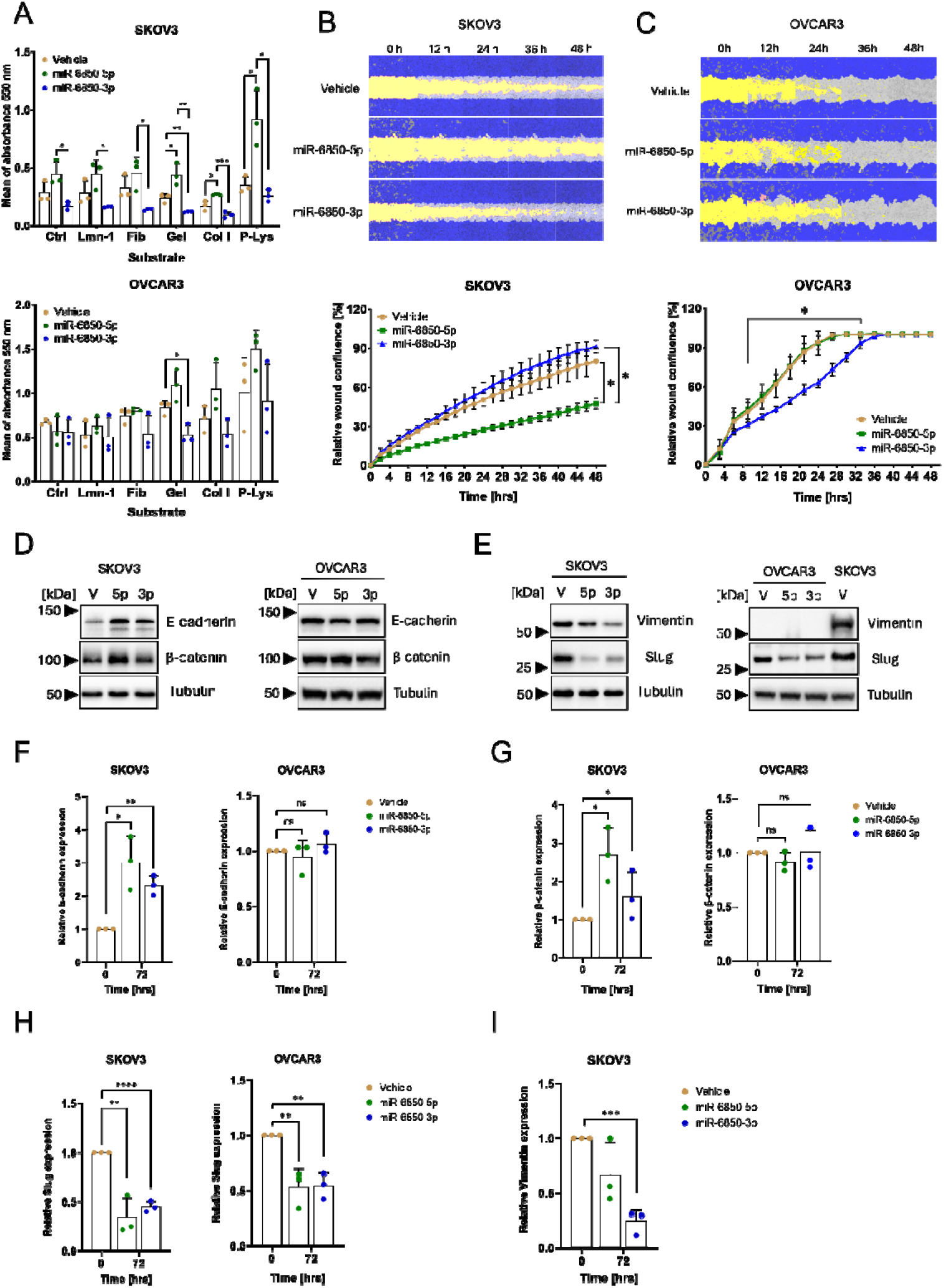
Functional impact of miR-6850 on cell adhesion, and migration. (A) Adhesion potential of SKOV-3 and NIH:OVCAR3 cells overexpressing miR-6850-5p, miR-6850-3p, and control cells. The adhesion potential was performed during the third-day post-transfection with miRNA mimics, on the following substrates: laminin-1 (Lmn-1), fibronectin (Fbn), gelatin (Gel), collagen I (Col 1), and poly-L-lysine (P-Lys). Scratch assay of (B) SKOV-3 and (C) NIH:OVCAR3 cells transfected with miR-6850-5p and miR-6850-3p, and control cells. Each panel contains the representative pictures of changes in the cell migration and graphs presenting a relative wound confluence reported in percentage, measured by Incucyte S3 at the time indicated above. The starting time (t = 0 h) indicates the SKOV-3 and NIH:OVCAR3 cells after three days post-transfection with miRNA mimics. The dark blue color represents the cells in confluence, the yellow color represents the performed wound while the light blue or grey color indicates the cells that migrated to the wound. Representative blots for (D) E-cadherin, β-catenin and (E) Slug, Vimentin in SKOV-3 (left) and NIH:OVCAR3 (right). Tubulin was used as a loading control. V – vehicle; 5p – miR-6850-5p; 3p – miR-6850-3p. The numbers on the left side of immunoblots represent the closest protein marker expressed in kDa. Densitometric quantification of (F) E-cadherin, (G) β-catenin, (H) Slug, and (I) Vimentin protein level in SKOV-3 and NIH:OVCAR3 cell lines from panels D and E. All experiments were conducted in three biological replicates and all data are shown as mean±SD. *p<0.05; **p<0.01; ***p<0.001, ****p<0.0001 determined by unpaired Student’s t-test.

In the next step, we assessed the migratory capacity of SKOV-3 and NIH:OVCAR3 cells using a scratch assay, initiated three days post-transfection. Relative wound confluence was measured at regular intervals over 48 hours. SKOV-3 cells transfected with miR-6850-5p displayed reduced wound closure compared to both control cells and miR-6850-3p-transfected cells. Interestingly, miR-6850-3p-transfected SKOV-3 cells exhibited a slight increase in migratory activity compared to control cells (Fig.3B). In contrast, NIH:OVCAR3 cells overexpressing miR-6850-3p displayed reduced wound closure while no difference was detected between control and miR-6850-5p overexpressing NIH:OVCAR3 (Fig.3C).

Since the phenotypic data pointed towards a modulation of cell cycle, proliferation, adhesion and migration by miR-6850, we reasoned that it may be involved in the regulation of EMT in HGSOC. To address this point, we conducted a comprehensive analysis of key EMT-associated protein markers. Specifically, we focused on epithelial markers, including E-cadherin and β-catenin, as well as mesenchymal markers such as Slug and Vimentin. These markers were selected due to their well-established involvement in the molecular reprogramming that underlies EMT, a critical event in tumor progression and metastasis [31].

In SKOV-3 cells overexpressing either miR-6850-5p or miR-6850-3p, we observed a clear shift toward an epithelial phenotype, characterized by increased expression of epithelial markers at the protein level (Fig.3D left). Specifically, E-cadherin levels increased by approximately threefold, while β-catenin levels rose up to twofold compared to control-transfected cells (Fig.3F and 3G). In contrast, NIH:OVCAR3 cells transfected with the same miRNA mimics did not show any appreciable changes in the expression of these markers (Fig.3D right, 3F and 3G), indicating a cell line-dependent response to miR-6850 overexpression. Concomitantly, SKOV-3 cells exhibited a strong reduction in mesenchymal markers (Fig.3E left). Slug, a key EMT-inducing transcription factor, was downregulated by up to fourfold, and Vimentin, a mesenchymal filament protein associated with motility [32] decreased by as much as sevenfold upon transfection with either miR-6850-5p or miR-6850-3p (Fig. 3H, 3I). In NIH:OVCAR3 cells, Slug levels were reduced by approximately twofold (Fig.3H), while Vimentin remained undetectable (Fig.3E right).

These results collectively suggest that miR-6850-5p and miR-6850-3p influence key traits linked to cancer progression, including cell adhesion, migration and may suppress EMT-evident through upregulation of epithelial markers and downregulation of mesenchymal transcription factor in cells where EMT is active.

### 3.5 miR-6850 regulates translation and the PI3K/Akt/mTOR pathway in HGSOC cells

The PI3K/Akt/mTOR pathway is frequently deregulated in HGSOC [33], with approximately 70% of OC cases exhibiting its aberrant activation, thereby promoting cell growth, proliferation, and cell survival through an intricate series of hyperactive signaling cascades [34]. Notably, SKOV-3 cells harbor an activating mutation in the *PIK3CA* gene, which is expected to result in constitutive activation of PI3K/Akt/mTOR axis [30]. In contrast, NIH:OVCAR3 cells are wild-type for *PIK3CA* and thus serve as a useful comparator model. This molecular distinction provided a rationale for investigating whether miR-6850 modulates PI3K signaling both in the presence and absence of pathway activation caused by genetic mutations.. To assess whether miR-6850 influences this pathway, we analyzed the impact of its overexpression on key signaling components, including phosphorylation of Akt kinase, eS6, and 4E-BP1, as well as expression of PTEN, a critical negative regulator of the PI3K pathway (27).

Our results demonstrated that overexpression of both miR-6850-5p and miR-6850-3p led to reduced phosphorylation levels of Akt, eS6, and 4E-BP1 in SKOV-3 and NIH:OVCAR3 cells, compared with the control (Fig. 5A). Notably, PTEN expression was upregulated approximately 2.5-fold following transfection with either miR-6850-5p or miR-6850-3p in SKOV-3 cells, while no significant change was observed in NIH:OVCAR3 cells (Fig. 4A and B). Phosphorylation of Akt was dramatically reduced—up to 12.5-fold in SKOV-3 and 2.8-fold in NIH:OVCAR3 cells (Fig. 5C). Interestingly, both miR-6850-5p and miR-6850-3p had similar effects on Akt phosphorylation in SKOV-3 cells, while in NIH:OVCAR3, miR-6850-5p induced a more pronounced inhibitory effect (Fig. 5C). eS6 phosphorylation was decreased by up to 2.5-fold in SKOV-3 and 3-fold in NIH:OVCAR3 cells, with miR-6850-3p showing the strongest effect in both cell lines (Fig. 5D). Additionally, both miR-6850-5p and miR-6850-3p overexpression in SKOV-3 and NIH:OVCAR3 led to a loss of the hyperphosphorylated 4E-BP1 band observed in vehicle, with a shift toward phosphorylated and non-phosphorylated forms, indicating inhibition of mTOR-dependent 4E-BP1 phosphorylation (Fig.5A)

**Fig. 5.**
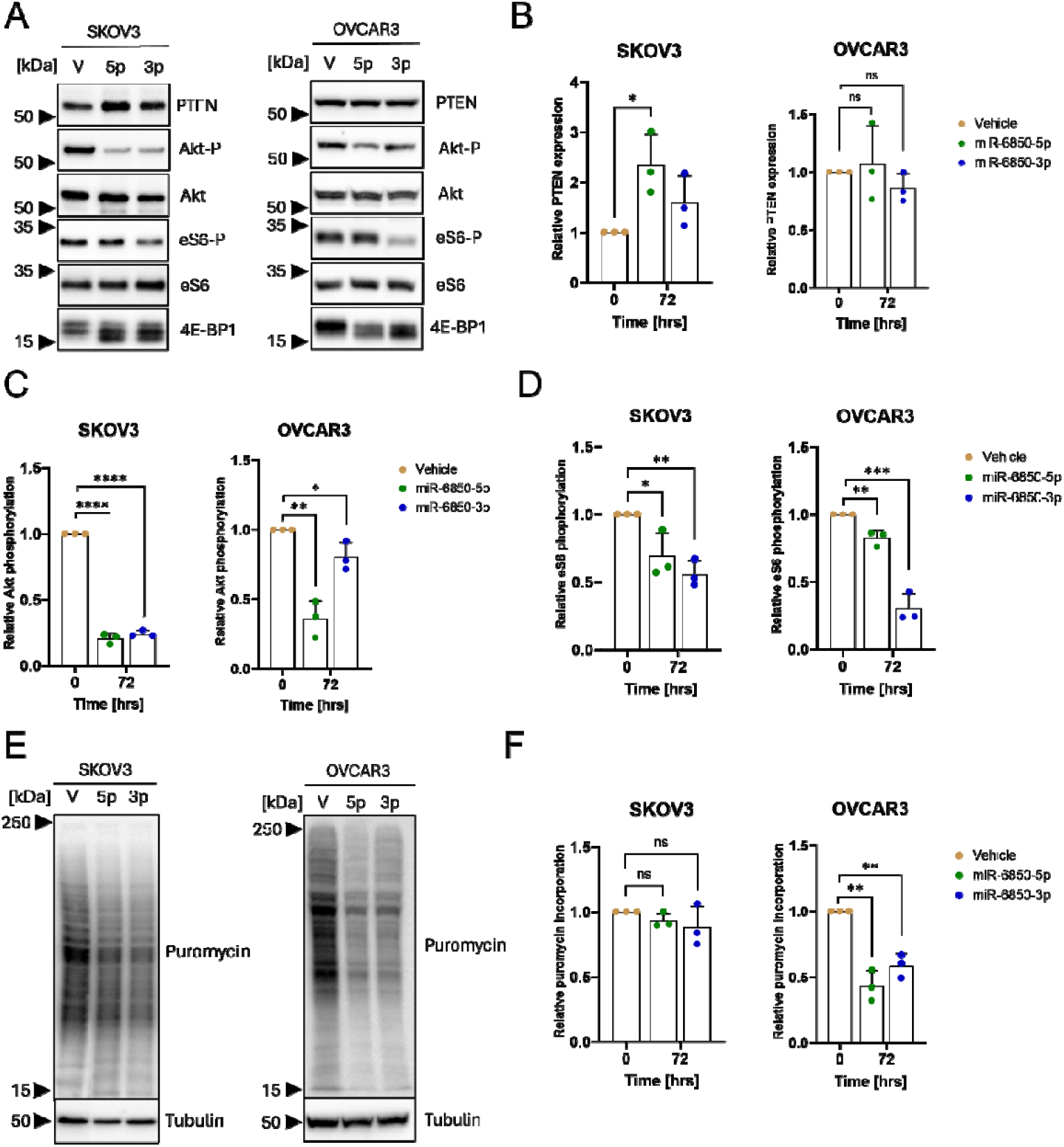
Effect of miR-6850 on PI3K/Akt/mTOR pathway and protein synthesis. (A) Representative blots for phosphorylated Akt, eS6, and 4E-BP1 and total PTEN in SKOV-3 (left) and NIH:OVCAR3 (right). Akt and eS6 total were used as loading controls. Densitometric quantification of (B) PTEN, (C) Akt phosphorylation, and (D) eS6 phosphorylation from panel A. (E) Representative blot of puromycin incorporation in SKOV-3 (left) and NIH:OVCAR3 (right). Tubulin was used as a loading control. V – vehicle; 5p – miR-6850-5p; 3p – miR-6850-3p. The numbers on the left side of immunoblots represent the closest protein marker expressed in kDa. (F) Densitometric quantification of translation efficiency from panel E. All experiments were conducted in three biological replicates and all data are shown as mean ± SD. *p<0.05; **p<0.01; ***p<0.001, ****p<0.0001 determined by unpaired Student’s t-test.

Given the involvement of miR-6850 in modulating the PI3K/Akt/mTOR pathway, we next assessed its impact on global protein synthesis using puromycin incorporation assays. Overexpression of miR-6850-5p and miR-6850-3p led to a modest reduction in puromycin-labeled proteins in SKOV-3 cells, whereas a more pronounced decrease was observed in NIH:OVCAR3 cells (Fig. 5E). Densitometric analysis of the puromycin signal revealed a ∼1.13-fold decrease in SKOV-3 cells upon miR-6850 overexpression (Fig. 5F). In NIH:OVCAR3 cells, miR-6850-3p and miR-6850-5p overexpression resulted in approximately 2-fold and 2.5-fold reductions in global protein synthesis, respectively (Fig. 5F), highlighting significant differences of miR-6850 on translational output based on the cellular context.

Collectively, these findings indicate that miR-6850 modulates the PI3K/Akt/mTOR signaling axis, leading to downregulation of protein synthesis in HGSOC models, and suggest a potential regulatory role of miR-6850 in controlling oncogenic translational programs.

### 3.6 miR-6850 regulates a distinct set of target genes

To gain insight into the molecular effectors through which miR-6850 may exert its tumor-suppressive functions, we performed an in silico target prediction analysis using six independent databases: miRPathDB, DIANA-microT, miRTarBase, miRWalk, miRDB, and TargetScan. The intersection of these datasets revealed a restricted number of overlapping targets (5 and 8 for miR-6850-3p and miR-6850-5p, respectively), suggesting a high degree of selectivity (Fig. 6A, B). Among the candidates selected for experimental validation, miR-6850-3p was predicted to regulate *PITX1, MBOAT1, TMBIM4, POLH*, and *SERTAD3*, while miR-6850-5p was predicted to target *DUT, MICAL2, OXTR, PLEKHA1, RUNX1, TP53INP1, XIAP*, and *ZNF446*. Functionally, the predicted targets of miR-6850 are involved in diverse cellular processes, including transcriptional regulation (*PITX1* [36], *SERTAD3* [37], *RUNX1* [38], *ZNF446* [39]), DNA repair and replication (*POLH* [40], *DUT* [41]), apoptosis and stress response (*XIAP* [42], *TP53INP1* [43], *TMBIM4* [44]), cytoskeletal dynamics (*MICAL2* [45], *PLEKHA1* [46]), and membrane signaling (*OXTR* [47], *MBOAT1* [48]). To experimentally assess the effect of miR-6850 on gene expression, we evaluated the mRNA levels of selected predicted targets in SKOV-3 and NIH:OVCAR3 cells following transfection with miR-6850 mimics. qRT-PCR analyses demonstrated that most of these targets were significantly regulated upon miR-6850 overexpression in both SKOV-3 and NIH:OVCAR3 cells, with the exception for *OXTR* (Fig. 6C, D). Notably, *TMBIM4* and *TP53INP1* transcripts were upregulated following transfection with miR-6850-3p and miR-6850-5p, respectively.

**Fig. 6.**
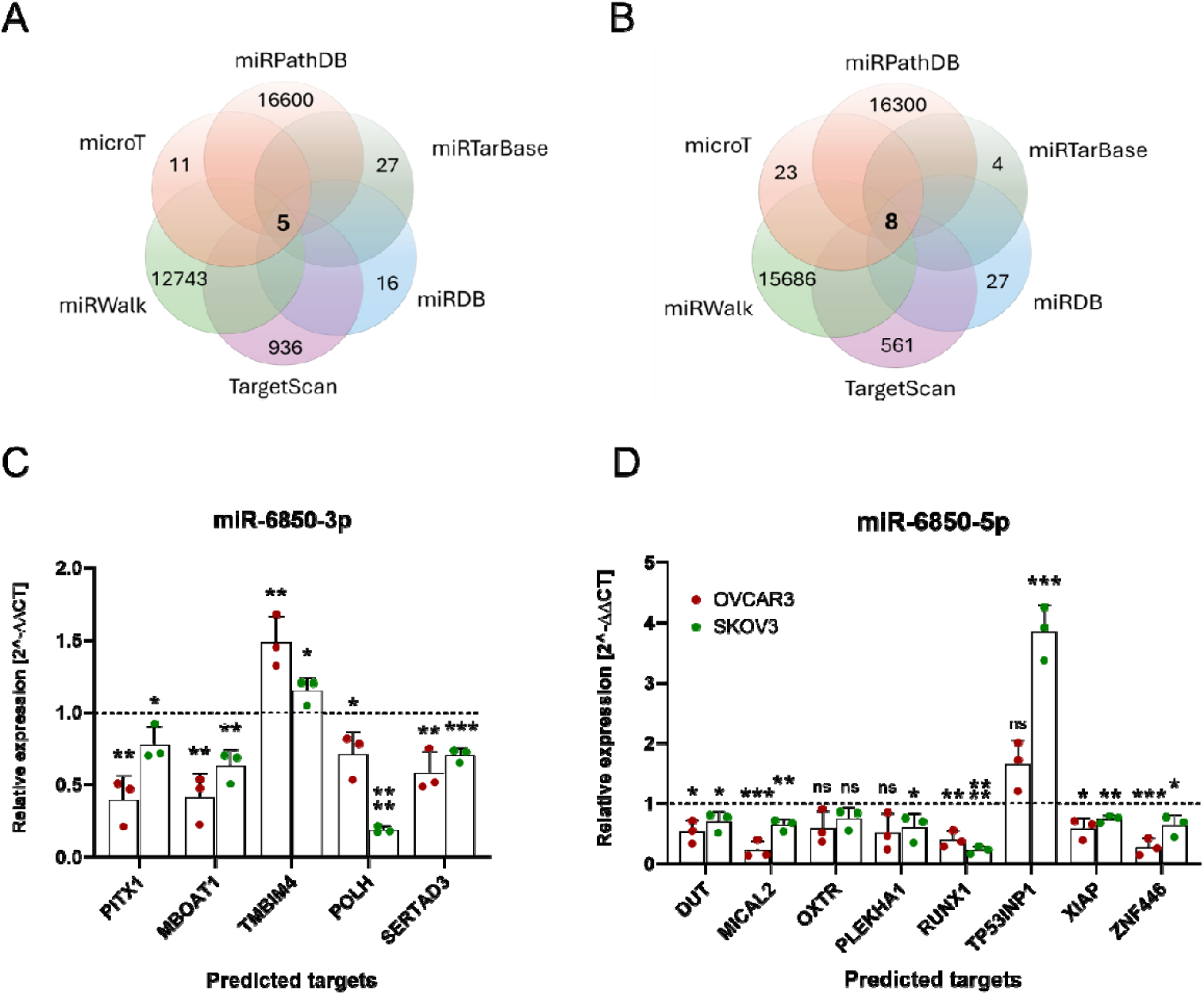
Prediction and validation of potential miR-6850 targets. (A–B) Venn diagrams illustrating the overlap of predicted target genes for (A) miR-6850-3p and (B) miR-6850-5p obtained from miRPathDB, DIANA-microT, miRTarBase, miRWalk, miRDB, and TargetScan databases. Common genes identified across all databases are indicated in the center of each diagram. (C–D) Relative mRNA expression levels of selected predicted targets in SKOV3 and NIH:OVCAR3 cells transfected with miR-6850-3p (C) or miR-6850-5p (D) compared to control cells, measured by qRT-PCR. All experiments were conducted in three biological replicates and all data are shown as mean ± SD. *p<0.05; **p<0.01; ***p<0.001, ****p<0.0001 determined by unpaired Student’s t-test.

Taken together, these findings suggest that miR-6850 exerts its effects through a multilayered regulatory network integrating transcriptional control, DNA repair, and apoptotic signaling, reinforcing its role as a context-dependent tumor suppressor in HGSOC.

## 4. Discussion

Ovarian cancer is a major global health problem. The high mortality rate has not improved for decades due to a lack of screening programs that could detect this cancer at an early stage and the poor efficacy of antitumor therapies in the long term [3]. Despite extensive research efforts, appropriate and effective markers for diagnostic purposes and targeted therapeutic approaches are still lacking. Recent studies indicate that miRNAs can serve as hallmarks for cancer diagnosis [49, 50] and/or cancer biomarkers [49, 51]. Here, we shed light on the critical role of miR-6850 in modulating the progression of HGSOC. Our initial bioinformatic analyses revealed that *MIR6850* is frequently amplified across multiple human cancers, yet the highest amplification rate was observed in HGSOC. MIR6850 is located in 8q24.3 locus, a region known for genomic instability in ovarian cancer [26, 27]. Despite this amplification, miR-6850 was among the least expressed miRNAs encoded at the 8q24.3 locus in both HGSOC tumor samples and OC cell lines, suggesting that amplification alone does not drive its expression. This discrepancy may stem from epigenetic silencing mechanisms, as *MIR6850* lies within a CpG island, a genomic context commonly associated with DNA methylation-mediated transcriptional repression [52]. Notably, other colocalized miRNAs within 8q24.3, such as miR-937 and miR-939, displayed much higher expression across both datasets, emphasizing the selective underexpression of miR-6850 despite its frequent amplification. In addition, our analyses revealed a consistent strand bias in miR-6850 expression, with miR-6850-5p being more abundant than miR-6850-3p in both tumor samples and OC cell lines. The preferentaial expression of the 5p strand could result from asymmetric strand selection during RISC loading, enhanced 5p strand stability, or differential degradation of the 3p strand. This strand bias is a well-recognized feature of miRNA biogenesis and function, often reflecting the dominant regulatory activity of one strand over the other [53]. Interestingly, although *MIR6850* is embedded within the first intron of the *RPL8* gene, its expression only weakly correlated with RPL8 mRNA levels. This suggests that miR-6850 may be regulated independently from its host gene, a phenomenon previously described for other intronic miRNAs [54].

Clinically, we observed that MIR6850 genomic alternation was associated with improved disease-specific survival in HGSOC patients. Furthermore, in a recent study, it has been shown that a decreased level of miR-6850-5p in serum has been detected in patients with benign ovarian cysts when compared to patients with epithelial ovarian cancer and healthy individuals [13]. In contrast, higher levels of miR-6850-5p in serum were associated with poorer progression-free survival in OC patients [55]. These examples illustrate that miR-6850 may be an emerging biomarker for OC patients.

To explore the functional relevance of miR-6850, we firstly examined the basal levels of miR-6850-5p and miR-6850-3p across OC cell lines, as compared to ovarian surface epithelial cells (HOSE) as healthy control. Indeed, ovarian surface epithelial cells may be the cell of origin of ovarian serous carcinoma, together with fallopian tube surface epithelial cells [56]. Both strands were consistently low expressed in HGSOC cell lines compared to the HOSE cell line, with 5p strand expressed at higher levels than 3p strand, confirming our bioinformatic analyses. Among the OC cell lines tested, SKOV-3 exhibits the highest endogenous expression of miR-6850 and displays a partial EMT phenotype together with a gain-of-function mutation in *PIK3CA* [30] that drives constitutive activation of the PI3K/Akt/mTOR pathway.. Conversely, NIH:OVCAR3 cells retain a fully epithelial phenotype, wild-type *PIK3CA* and *PTEN* [30], and low miR-6850 levels. The use of these two phenotypically and molecularly distinct models enabled us to dissect miR-6850’s regulatory roles in both mesenchymal-like and epithelial contexts, providing insight into how its activity interfaces with EMT programs and PI3K/Akt/mTOR signaling dynamics. Notably, we demonstrated that re-expression of either of the two strands miR-6850-5p and miR-6850-3p into HGSOC cells suppressed tumorigenic behaviors. Both SKOV-3 and NIH:OVCAR3 cells exhibited reduced proliferation upon miR-6850 overexpression, however, the inhibitory effect was more pronounced in SKOV-3 cells, where both strands led to a marked and sustained growth reduction over time. In contrast, NIH:OVCAR3 cells miR-6850-5p exerting a stronger antiproliferative effect than miR-6850-3p. This difference suggests the functional impact of each strand is context-dependent and may relate to intrinsic differences in the baseline phenotype or signaling landscape of the two cell lines. Furthermore, NIH:OVCAR3 were more susceptible to the long-term growth-suppressive effects of miR-6850-5p, as shown by a grater reduction in colony number and area. Both isoforms significantly downregulated Cyclin A2 and Cyclin B1 expression. This supports the notion that miR-6850 exerts anti-proliferative effects by targeting components of the cell cycle machinery, in line with reports of other miRNAs such as miR-125b, miR-24 and let-7, which similarly repress cyclin expression to inhibit cell division [57–59].

A key highlight of our study is the observation that miR-6850 promotes a mesenchymal-to-epithelial transition (MET), particularly in SKOV-3 cells, which exhibit a more mesenchymal-like phenotype [29]. Upon miR-6850 overexpression, we observed a robust upregulation of epithelial markers (E-cadherin, β-catenin) and downregulation of mesenchymal markers (Slug, Vimentin). This shift toward an epithelial state suggests a reversal of EMT, a hallmark of metastatic potential in cancer. Consistently, the differential effects of miR-6850 on cell adhesion and migration reflect the interplay between integrin activity and epithelial differentiation. In SKOV-3 cells, which express high levels of α5β1 and αvβ3 integrins [60], miR-6850-5p enhanced adhesion to ECM components such as fibronectin and gelatin, yet concurrently reduced migration. This suggests that increased substrate binding may impair the dynamic adhesion turnover required for motility. Additionally, miR-6850-driven upregulation of E-cadherin likely stabilized cell–cell junctions and functionally suppressed integrin-mediated signaling, further contributing to reduced migration. In NIH:OVCAR3 cells, which already express high baseline levels of E-cadherin [29] and have reduced levels of these integrins [60], miR-6850 had a milder effect on adhesion but significantly impaired migration, particularly with the 3p strand. This may reflect reinforcement of epithelial traits and inhibition of Akt signaling, rather than major changes in integrin engagement. These findings underscore the importance of cellular context in shaping miRNA function and further suggest that miR-6850-5p may exert more potent anti-metastatic effects than miR-6850-3p, particularly in mesenchymal-like cell states. The ability of miR-6850 to modulate EMT has profound implications for metastasis suppression, and may offer a therapeutic route to prevent or reverse invasive behavior in HGSOC.

In addition to its role in phenotypic plasticity, miR-6850 also modulated intracellular signaling networks, specifically the PI3K/Akt/mTOR pathway, which is commonly hyperactivated in ovarian cancer [33]. We found that both miR-6850 isoforms suppressed Akt phosphorylation and downstream targets such as eS6 and 4E-BP1. Importantly, this was accompanied by upregulation of PTEN, a major negative regulator of PI3K signaling, in SKOV-3 cells. The observed cell line differences, such as the lack of PTEN modulation in NIH:OVCAR3, may reflect distinct mutational landscapes or feedback mechanisms. Nonetheless, the consistent downregulation of Akt and mTOR signaling across models indicates that miR-6850 plays a broader role in repressing oncogenic translation and metabolism. However, the impact on global protein synthesis appeared **cell-type specific**. Puromycin incorporation assays revealed only a modest reduction in SKOV-3 cells, whereas NIH:OVCAR3 cells showed a pronounced decrease, particularly upon miR-6850-5p overexpression. This discrepancy may reflect differences in baseline translational activity, dependence on mTOR signaling, or the relative contribution of miR-6850 targets in each context. Given the essential role of protein synthesis in sustaining malignant growth, the ability of miR-6850, especially in epithelial-like cells to suppress translation adds a further dimension to its tumor-suppressive function.

In line with the pleiotropic functions observed in our cellular assays, bioinformatic analyses identified multiple candidate targets of miR-6850 involved in stress response, DNA repair, cytoskeletal dynamic, and transcriptional regulation. The downregulation of most targets upon miR-6850 overexpression suggest that miR-6850 may predominantly act as a post-transcriptional repressor, likely through binding to the 3’ untranslated regions (UTRs) of target mRNAs. Among these, RUNX1 and MICAL2 have been shown to regulate survival, EMT programs, and proliferation in ovarian cancer, rescpectively [61–63]. Silencing of RUNX1 reduced EMT and increased apoptosis via the FOXO1-Bcl2 axis [62] while MICAL2 silencing suppressed ovarian cancer proliferation and migration, promoting a MET [63]. The observed reduced proliferation, migration and changes in EMT markers expression upon miR-6850-5p overexpression is consistent with RUNX1 and MICAL2 repression. Cellectively, these targets suggest that miR-6850-5p mediates tumor-suppressive effects in HGSOC through a coordinated inhibition of EMT process and cytoskeletal remodeling, acting as a central regulator of cellular plasticity and survival. Interestingly, two targets, namely TMBIM4 and TP53INP1, were upregulated following miR-6850 overexpression. This observation rises the possibility that miR-6850 may also interact with the 5’UTRs of these mRNAs, leading to increased transcript level. Notably, previous studies have shown that miR-10a [64] and miR-103a-3p [65] bind to the 5’UTR of ribosomal proteins and GPRC5A mRNAs, respectively, enhancing their translation and increased protein abundance.

Taken together, our study establishes miR-6850 as a multifaceted regulator of HGSOC progression, impacting cell proliferation, adhesion, migration, epithelial plasticity, and protein biosynthesis. These effects are likely mediated through the coordinated regulation of targets involved in transcriptional control, DNA replication, and cytoskeletal dynamics. Importantly, while both miR-6850-5p and miR-6850-3p are functionally active, miR-6850-5p consistently exerted stronger effects in phenotypic and biochemical assays, suggesting therapeutic prioritization of this strand.

Despite these insights, our study has limitations. The use of in vitro models, although informative, does not fully capture the complexity of the tumor microenvironment or immune interactions. Future work will need to assess miR-6850 function in more complex models, like 3D cultures and orthotopic animal models of ovarian cancer. Moreover, understanding whether miR-6850 modulates chemotherapy response, and identifying direct mRNA targets, could open new avenues for therapeutic development.

## 5. Conclusions

In conclusion, miR-6850-5p and miR-6850-3p act as potent tumor suppressors in HGSOC by modulating cell proliferation, migration, adhesion, EMT, and the PI3K/Akt/mTOR pathway. Their low expression in ovarian cancer suggests the potential of miR-6850 as a biomarker and therapeutic target. This study lays the groundwork for future research focused on translating these findings into clinical applications, including evaluating the druggability of miR-6850 or its downstream targets, as well as assessing the correlation between tumor and circulating miR-6850 levels with outcomes and therapy responses in patients with high-grade serous ovarian cancer..

## Supporting information

Supplemental tabel 1

## Author Contributions

All authors contributed to the conceptualization and research design. KF and MP designed the research process. KF performed the experiments. KF, MP analyzed the data. KF, GG, IK and MP wrote the original draft, and reviewed, and edited the paper. KF, DP, IK, GG, and MP read and approved the final version of the submitted manuscript.

## Acknowledgments

The authors gratefully acknowledge the networking efforts of the TRANSLACORE Action (CA21154) by COST (European Cooperation in Science and Technology), which facilitated their professional connection, Dr. Dorian Forte for support with cell cycle analysis, and Prof. Lorenzo Montanaro and his research group for continuos discussion on the project.

## Availability of data

The data used to support the findings of this scientific work are available from the corresponding author upon reasonable request.

