## Supplemental tabel 1 for "miR-6850 Drives Phenotypic Changes and Signaling in High Grade Serous Ovarian Cancer"

| Antibody | Catalog number | Manufacturer | Dilution |
| --- | --- | --- | --- |
| anti-alpha-tubulin [DM1A] | Ab7291 | Abcam | 1:5000 |
| anti-puromycin | MABE343 | Millipore | 1:5000 |
| Anti-phospho-RPS6 (Ser235) | Sc-101793 | Santa Cruz Biotechnology | 1:1000 |
| Anti-RPS6 | Sc-74459 | Santa Cruz Biotechnology | 1:500 |
| Anti-phospho-Akt (Ser473) | #9271 | Cell Signaling Technology | 1:1000 |
| Anti-Akt | #9272 | Cell Signaling Technology | 1:1000 |
| Anti-E-cadherin | #3195 | Cell Signaling Technology | 1:1000 |
| Anti-Slug | #9585 | Cell Signaling Technology | 1:1000 |
| Anti-4E-BP1 | #9644 | Cell Signaling Technology | 1:1000 |
| Anti-Cyclin B1 | #4135 | Cell Signaling Technology | 1:1000 |
| Anti-Vimentin | #5741 | Cell Signaling Technology | 1:1000 |
| Anti-GAPDH | #2118 | Cell Signaling Technology | 1:1000 |
| Anti-PTEN | NCL-PTEN | Novocastra | 1:1000 |
| Anti-β-catenin | NCL-B-CAT | Novocastra | 1:1000 |
| Anti-Cyclin A2 | NCL-CYCLIN A | Novocastra | 1:500 |
| Peroxidase IgG Fraction Monoclonal Mouse Anti-Rabbit IgG, light chain specific | 211-032-171 | Jackson ImmunoResearch | 1:5000 |
| Peroxidase AffiniPure Goat Anti-Mouse IgG, light chain specific | 115-035-174 | Jackson ImmunoResearch | 1:5000 |
